# Rapid Paediatric Sequencing (RaPS): Comprehensive real-life workflow for rapid diagnosis of critically ill children

**DOI:** 10.1101/283697

**Authors:** Lamia Boukhibar, Emma Clement, Wendy Jones, Suzanne Drury, Louise Ocaka, Andrey Gagunashvili, Polona Le Quesne Stabej, Chiara Bacchelli, Nital Jani, Shamima Rahman, Lucy Jenkins, Jane Hurst, Maria Bitner-Glindzicz, Mark Peters, Philip Beales, Hywel J Williams

## Abstract

**Background:** Rare genetic conditions are frequent risk factors for, or direct causes of, organ failure requiring paediatric intensive care unit (PICU) support. Such conditions are frequently suspected but unidentified at PICU admission. Compassionate and effective care is greatly assisted by definitive diagnostic information. There is therefore a need to provide a rapid genetic diagnosis to inform clinical management.

To date, Whole Genome Sequencing (WGS) approaches have proved successful in diagnosing a proportion of children with rare diseases, but results may take months to report or require the use of equipment and practices not compatible with a clinical diagnostic setting. We describe an end-to-end workflow for the use of rapid WGS for diagnosis in critically ill children in a UK National Health Service (NHS) diagnostic setting.

**Methods:** We sought to establish a multidisciplinary Rapid Paediatric Sequencing (RaPS) team for case selection, trio WGS, a rapid bioinformatics pipeline for sequence analysis and a phased analysis and reporting system to prioritise genes with a high likelihood of being causal. Our workflow was iteratively developed prospectively during the analysis of the first 10 children and applied to the following 14 to assess its utility.

**Findings:** Trio WGS in 24 critically ill children led to a molecular diagnosis in ten (42%) through the identification of causative genetic variants. In three of these ten individuals (30%) the diagnostic result had an immediate impact on the individual’s clinical management. For the last 14 trios, the shortest time taken to reach a provisional diagnosis was four days (median 7 days).

**Interpretation:** Rapid WGS can be used to diagnose and inform management of critically ill children using widely available off the shelf products within the constraints of an NHS clinical diagnostic setting. We provide a robust workflow that will inform and facilitate the rollout of rapid genome sequencing in the NHS and other healthcare systems globally.

**Funding:** The study was funded by NIHR GOSH/UCL BRC: ormbrc-2012-1

## Introduction

An increasing proportion of critically ill children have one or more chronic diseases that contribute to, or directly precipitate, paediatric intensive care admission (1). Rare genetic conditions are present in a significant proportion of elective and emergency admissions. Uncertainty about diagnosis and often prognosis contributes to the difficulty of planning optimal care. Achieving a rapid molecular diagnosis in critically ill children with a rare genetic disease may improve the basis for such plans including informing on the potential value of highly invasive treatments (2, 3). Reaching a genetic diagnosis also precludes the need for further diagnostic investigations, which may be invasive, painful, and expensive (4). For the family, a molecular diagnosis enables accurate genetic counselling and ends the diagnostic odyssey (5). However, obtaining a genetic diagnosis in a timely manner in critically ill individuals is frequently challenging and often not possible. Factors preventing a rapid genetic diagnosis include heterogeneity of disease, limited availability of broad genetic testing, long time frames involved in standard diagnostic molecular testing, and limited knowledge of the molecular basis for most genetic disorders.

Recent advances in genome sequencing and bioinformatics provide a solution to many of the traditional hurdles presented by rare diseases. Whole exome sequencing (WES) approaches, where only the coding sequence of genes is targeted, have proven successful in diagnosing a proportion of children and adults with rare diseases in both the research and diagnostic arenas (5-9). Whole genome sequencing (WGS), which unlike WES is not biased to particular genomic regions, is now frequently being performed in the research setting and is beginning to be used for diagnostic purposes. A comparison between the two methods has shown that WGS is the preferred option for testing Mendelian disorders (10). In the UK WGS is being extended into the healthcare environment through the 100,000 Genomes Project (11, 12), however, feedback of results is currently expected to take many months. Early studies in the USA (13, 14) and Netherlands (15) have shown a benefit for rapid sequencing in acutely ill children but these studies have used modified laboratory equipment and working procedures incompatible with standard diagnostic laboratory practices or have analysed a pre-determined gene list, respectively.

For this study, rapidity of diagnosis is not the only or even most important issue to address as sustainability of a rapid genome sequencing service in the context of a NHS diagnostic laboratory is of paramount importance. The aim of this study was therefore to expand on previous rapid sequencing studies by developing the first end-to-end workflow using rapid WGS to diagnose critically ill children in an NHS setting. Specifically, this begins with the identification of an eligible patient on the paediatric intensive care unit (PICU) and ends with the delivery of a diagnostic report. To do this we set up a multidisciplinary team to ensure our workflow seamlessly transitioned between the various specialities. We adopted a fully prospective two-stage approach whereby the first 10 trios were used to iteratively develop a workflow which was then applied to the next 14 trios.

An essential goal of this study was to develop a workflow integrated within an existing service laboratory that could be adopted by other diagnostic centres. To achieve this we only used off-the-shelf products and equipment and designed our protocol to fit with standard working practices. We make this information freely available for others to use.

## Methods

The study was undertaken in a UK National Health Service (NHS) tertiary children’s hospital with a 23-bedded multi-disciplinary PICU and a 20 bed paediatric cardiac intensive care unit.

The study has UK Research Ethics Committee approval REC reference 08/H0713/82. Signed informed parental consent for participation in this study was obtained in all cases.

### RaPid Sequencing (RaPS) Team

We established a multidisciplinary RaPS team consisting of clinical geneticists, research and clinical scientists. The RaPS team was supported by paediatric intensive care unit (PICU) clinicians and other paediatric specialist teams who identified critically ill individuals for inclusion. Our workflow comprises detailed inclusion and exclusion criteria, clinical data capture and conversion to Human Phenotype Ontology (HPO) terms, rapid DNA extraction and WGS, a rapid bioinformatics analysis pipeline, tiered reporting of potentially causative variants, multidisciplinary team discussion, and validation of results in an accredited NHS diagnostic laboratory (Figure 1).

**Figure 1.**
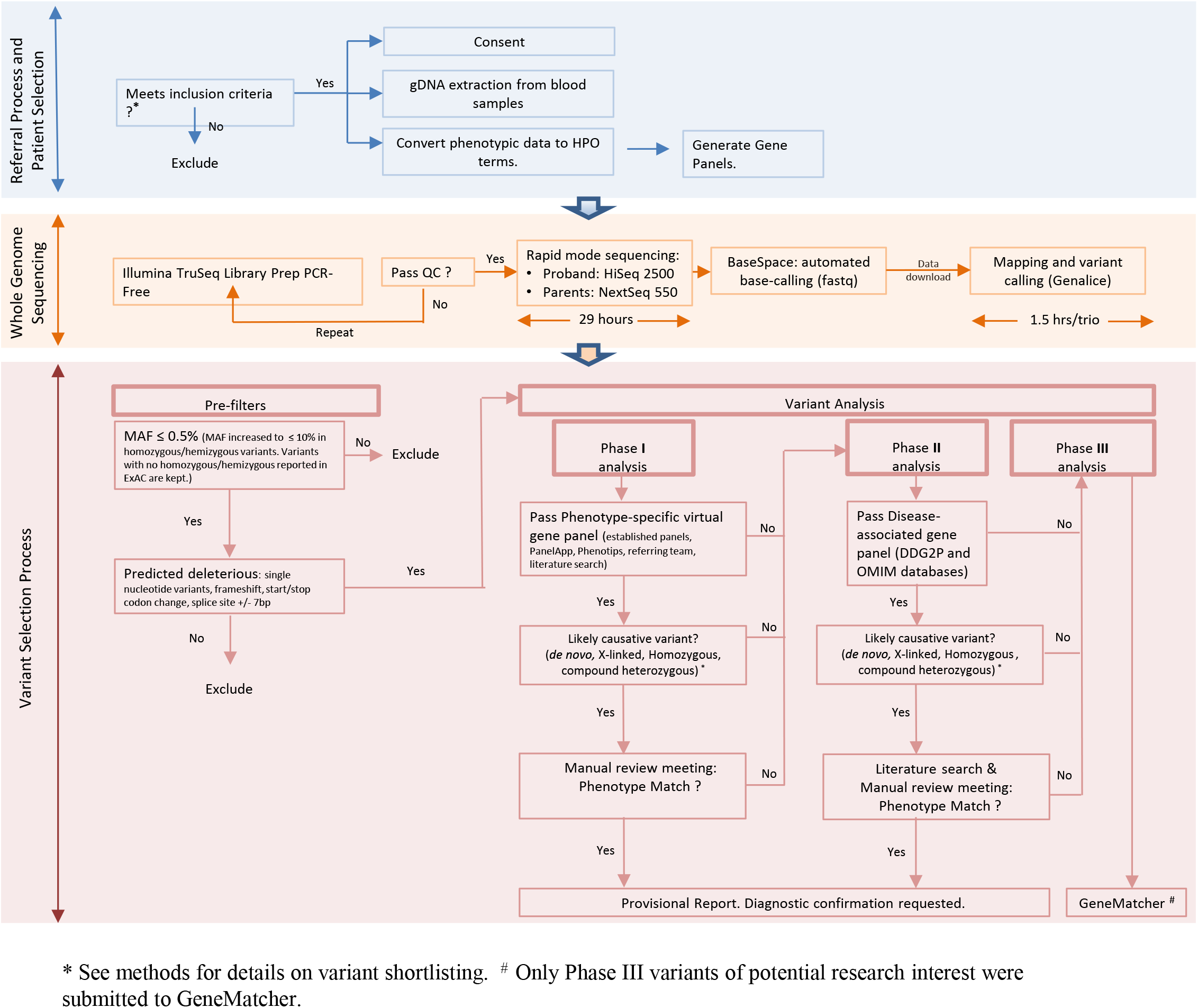
Description of RaPS workflow.

### Inclusion criteria

Suitable participants were clinically ascertained by a specialist physician or PICU consultant between August 2015 until October 2017. The first 10 trios were run as proof-of-principle to establish workflow systems. This allowed us to iteratively review our inclusion criteria and led to the development of the following list that we applied to the remaining 14 trios:

1. Essential inclusion criteria:
  a. Trio DNA samples must be available.
  b. Parental consent.
  c. Suspected underlying monogenic cause.
2. High priority criteria:
  a. A genetic diagnosis may significantly alter the clinical management of the patient.
  b. The phenotype or family history data strongly implicate a genetic aetiology, but the phenotype does not correspond with a specific disorder for which a genetic test targeting a specific gene is available on a clinical basis.
  c. A patient presents with a defined genetic disorder that demonstrates a high degree of genetic heterogeneity, making WGS analysis a more practical approach than a Gene Panel test.
  d. A patient presents with a likely genetic disorder, but specific diagnostic tests available for that phenotype have failed to arrive at a diagnosis or are not accessible within a reasonable timeframe. Such tests may include a Gene Panel test, Microarray, Biochemical test, Imaging or Biopsy.
  e. Imminent demise of patient not likely.

Using these criteria we aimed to identify individuals with a high likelihood of having a monogenic disorder and focused on those individuals in whom achieving a genetic diagnosis would be likely to inform clinical management in the acute setting. We also excluded individuals who were felt to be at high risk of imminent demise. For this group of children standard diagnostic testing or access to WGS through the 100,000 Genomes Project (100KGP) (7) was suggested where appropriate in order to offer an explanation and inform future genetic counselling for their parents and family members.

### Recruitment and Consent

To expedite the identification and pathogenic assessment of causative genes we recruited biological trios consisting of proband and both parents (14). A template for recording clinical and family history at time of consent was developed in order to standardise data capture and improve workflow. Phenotypic information provided by a clinical geneticist, or specialist paediatrician was captured as HPO terms to facilitate bespoke gene panel design for each patient (Supplementary Table 1).

Participants were given the choice of opting in or out of return of secondary findings as guided by recommendations from the American College of Medical Genetics and Genomics (ACMG)(16).

### Genomic Assays

Detailed methods are found in Supplementary Material 2. Briefly, DNA was extracted using Chemagic-STAR (Hamilton USA). Whole genome gDNA libraries were prepared using TruSeq DNA PCR-Free Library Prep (Illumina, USA) following manufacturer’s advice starting with 1ug of sheared gDNA. Parental samples were pooled at equimolar concentrations and sequenced on Illumina NextSeq 550 High-Output Mode (29 hours). Patient samples were sequenced on Illumina HighSeq 2500 Dual Flow Cell, Rapid Run Mode (27 hours) except for patient samples from the last two trios which were sequenced on NextSeq 550 High-Throughput Mode (29 hours). Mapping and variant calling were performed using a GENALICE appliance running GENALICE Map 2.5.5 including Mapping, Variant Calling and the Population Calling module for trio analysis (GENALICE BV, Netherlands). GENALICE default configuration files were used for WGS mapping, and trio variant detection.

### Variant interpretation

Ingenuity Variant Analysis™ software (QIAGEN, USA) was used to identify rare variants predicted to result in loss of function, or to have a functional effect on the protein. Variants with a frequency of ≤ 0·5% in 1000 Genomes (17), ExAC (18), and Exome Variant Server were investigated. Additionally, we also performed a complimentary analysis to identify candidate recessive and X-linked variants with a high carrier frequency. For this analysis we used a more permissive allele frequency cut off of ≤ 10% with those variants with no reported homozygous or hemizygous genotypes in ExAC included for further analysis. We then selected only variants that were predicted to be deleterious (simple nucleotide variants, frameshift, start/stop codon change, splice site +/- 7bp). Genetic filters were set to investigate autosomal recessive homozygous, autosomal recessive compound heterozygous, X-linked and de novo variants. For variants within genes with a recessive mode of inheritance in the Phase I analysis, we also shortlisted predicted deleterious heterozygous variants. When such variants were identified we manually inspected the genomic region using Integrative Genome Viewer (IGV) software (19) to detect potential structural variants on the second allele.

Data were analysed in a three-stage process (Phase I-III) to prioritise likely causative genes and facilitate prompt return of results (Figure 1). All putative pathogenic calls were manually assessed using IGV to ensure they were true variants and not technical artefacts.

### Phase I analysis

In Phase I we restricted the genes analysed to those with a high probability of being implicated in the individual’s disorder. This required analysis of a bespoke Phase I gene panel generated from gene lists provided by the referring clinical teams in conjunction with an HPO derived panel utilising the following resources: Genomics England PanelApp (20), Phenotips (21), and Online Inheritance in Man (OMIM) Gene Map (22) (see Supplementary Material 3). If a Phase I variant was deemed to be causal and explain the entire phenotype, no further analysis of WGS was deemed necessary.

### Phase II analysis

Individuals entered Phase II analysis when no likely-causative variant was identified in Phase I analysis or when a Phase I candidate variant did not fully explain the reported phenotype. Phase II comprised a broad analysis of genes known to be associated with developmental disorders and disease more generally. Phase II involved analysis of genes from the Developmental Disorders Genotype-Phenotype (DDG2P) database (6) and OMIM Morbid genes (22).

### Phase III analysis

Individuals entered Phase III analysis if no causal variants were identified from Phase I or Phase II or their phenotype was not fully explained. The aim of Phase III was to open up the analysis to select variants in any gene with compelling evidence for causality based on the deleteriousness of the variant and either animal models, expression pattern, or *in silico* predictions. Where a genetic diagnosis was not achieved in Phase I or II, variants of potential research interest from Phase III were shared with the online portal GeneMatcher (23) to identify potential collaborators with variants in the same gene.

### Multi-disciplinary Review

Variants identified from Phase I and II analysis were triaged by the core RaPS team including a Clinical geneticist and research scientists. Any variants deemed to be potentially relevant to the individual’s phenotype were scored according to ACMG variant interpretation guidelines (24). These were then reviewed in a genomic multidisciplinary team (MDT) meeting comprising at least two clinical geneticists, the referring team when available and clinical and research scientists in order to determine a consensus on pathogenicity and the need for further investigations.

### Feedback of Results

Variants assessed as pathogenic or likely pathogenic and contributing to the individual’s phenotype following MDT discussion were fed back to referring clinicians. At this point a provisional research results report was generated. Diagnostic results were validated in an accredited laboratory using Sanger sequencing. If no likely pathogenic variants were identified after Phase II analysis, a ‘no primary findings’ research results report was issued to the referring clinical team detailing the analysis performed and plan for continued research analysis.

## Results

### RaPS workflow: from patient to variant

Individuals recruited were on average known to eight specialist medical teams in the hospital (Figure 2 and Supplementary Figure 1). Mean age of affected individuals at point of sequencing was 15·86 months (range 7 days-13y 2 months) with a median age of 2.5 months (Supplementary Figure 2).

**Figure 2.**
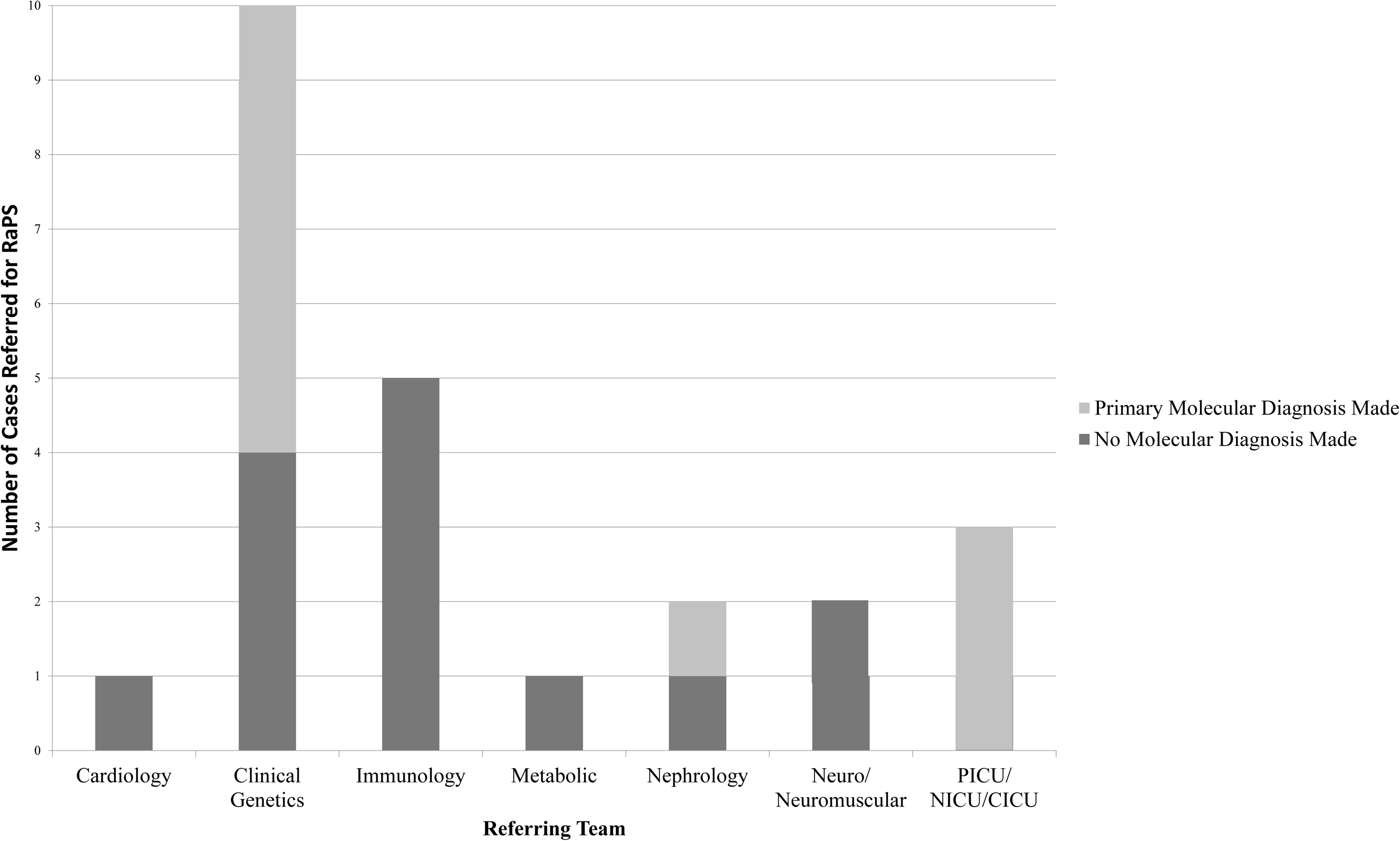
Number of Trios referred and Diagnoses made per Specialty.

Whole genome sequencing of the 24 trios generated an average of 5·8 million genomic variants per trio (Supplementary Table 2), including those seen in only one parent. The time taken for read mapping and variant calling of the sequence data using the GENALICE appliance ranged from 10 to 40 minutes with an average time of 19 minutes per sample. The number of variants per workflow stage and phase is indicated in Supplementary Figure 3. Our coverage metrics showed that on average 88% of the proband’s genome had at least 10X coverage and an average of 67% of the parent’s genome had at least 10X coverage (Supplementary Table 3). Similar coverage rates were obtained for the coding regions investigated during variant interpretation.

A primary molecular diagnosis (classified as a diagnosis accounting for the majority of an individual’s phenotype) was achieved in ten out of 24 trios (42%) (Table 1). Of note, all diagnoses were made in Phase I analysis (Supplementary Table 4). Diagnostic variants comprised four de novo mutations, three pairs of compound heterozygous variants and three homozygous variants.

**Table 1.**
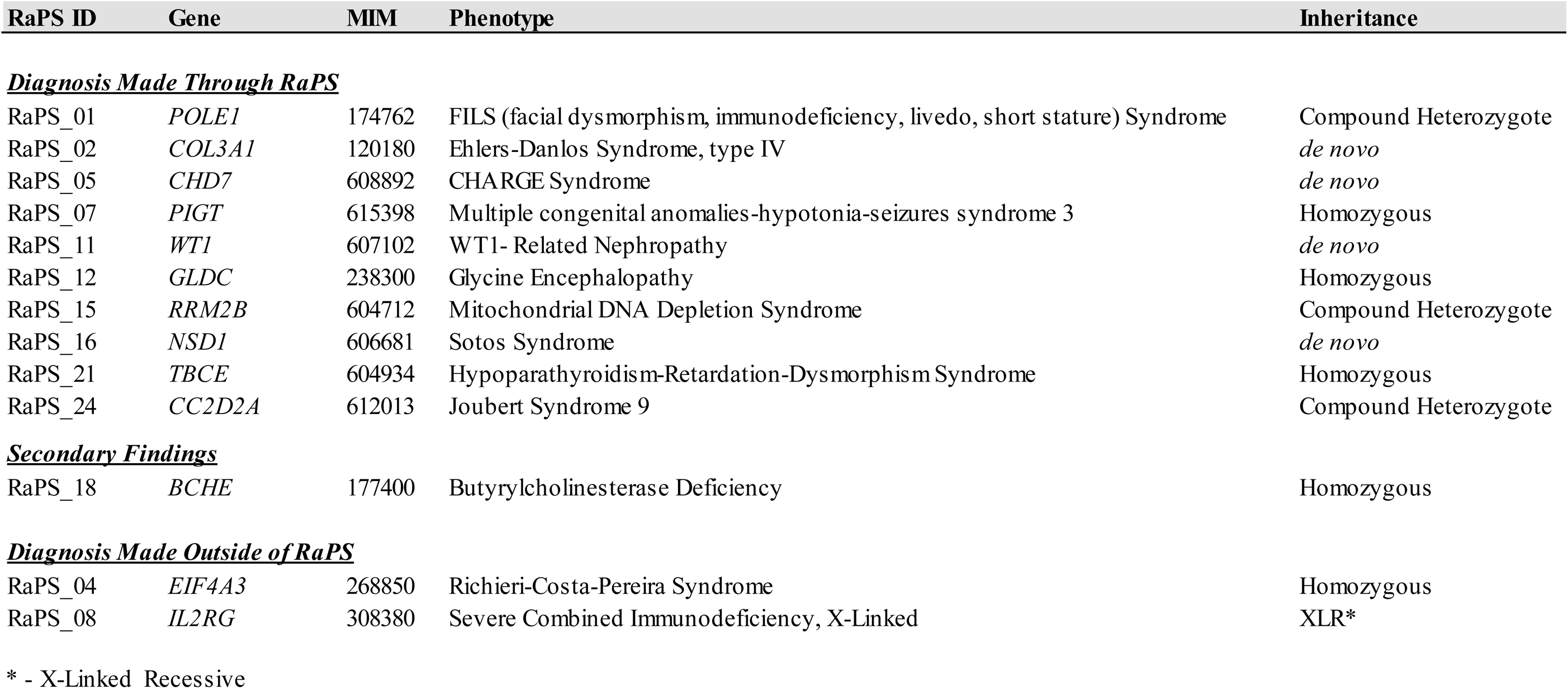
Summary of diagnoses made in the RaPS cohort. Ten diagnoses were made as a result of WGS through RaPS, all of which explain the primary clinical findings. In one case (RaPS_18) a secondary finding of homozygous *BCHE* mutations was identified and fed back to the referring team as it was deemed clincally relevant. Two molecular diagnoses were found outside of RaPS; A patient with a known mutation in *IL2RG* (RaPS_08) was recruited to RaPS to investigate dual pathology. The *IL2RG* mutation was confirmed but no second molecular diagnosis made. In RaPS_04, a homozygous 5’UTR expansion not detected by WGS was identified in *EIF4A3* by a different group.

In addition to the diagnostic variants identified in Phase I, a diagnostic 5’UTR expansion in *EIF4A3* was identified in an individual (RaPS_04) by a collaborative group (25) and not detected by our WGS analysis. In a further case we confirmed a previously identified *IL2RG* variant in a proband with immune deficiency. The referring team had requested RaPS analysis with a suspicion that there may be a second cause of the observed clinical features; however, no additional putatively causal variants were identified (Table 1).

In Phase II analysis a secondary finding was identified in one individual (RaPS_18). This comprised a homozygous variant in *BCHE.* Variants in *BCHE* are associated with post-anaesthetic apnoea (Table 1).

### Timelines for diagnosis

The shortest time taken to complete the full workflow (from consent to return of provisional diagnosis) was five days (RaPS_11). To allow a comparison to previous studies (13-15) we also measured the time to diagnosis for the last 14 trios from the point library preparation began to return of the provisional result. The shortest time for this time period was 4 days with a median time of 7 days (SD+/- 9.6 days) (Figure 3). This timeline reflects ‘real life’ and is based on the standard working hours of a diagnostic laboratory and includes technical delays caused by reagent failure or lack of availability of sequencers. The turnaround time for the first 10 ‘proof of principle’ cases were much more variable whilst systems and workflow was being established. Other factors that resulted in an increased time to diagnosis include delays before the library preparation started and included the non-availability of a parent for consent or blood draw.

**Figure 3:**
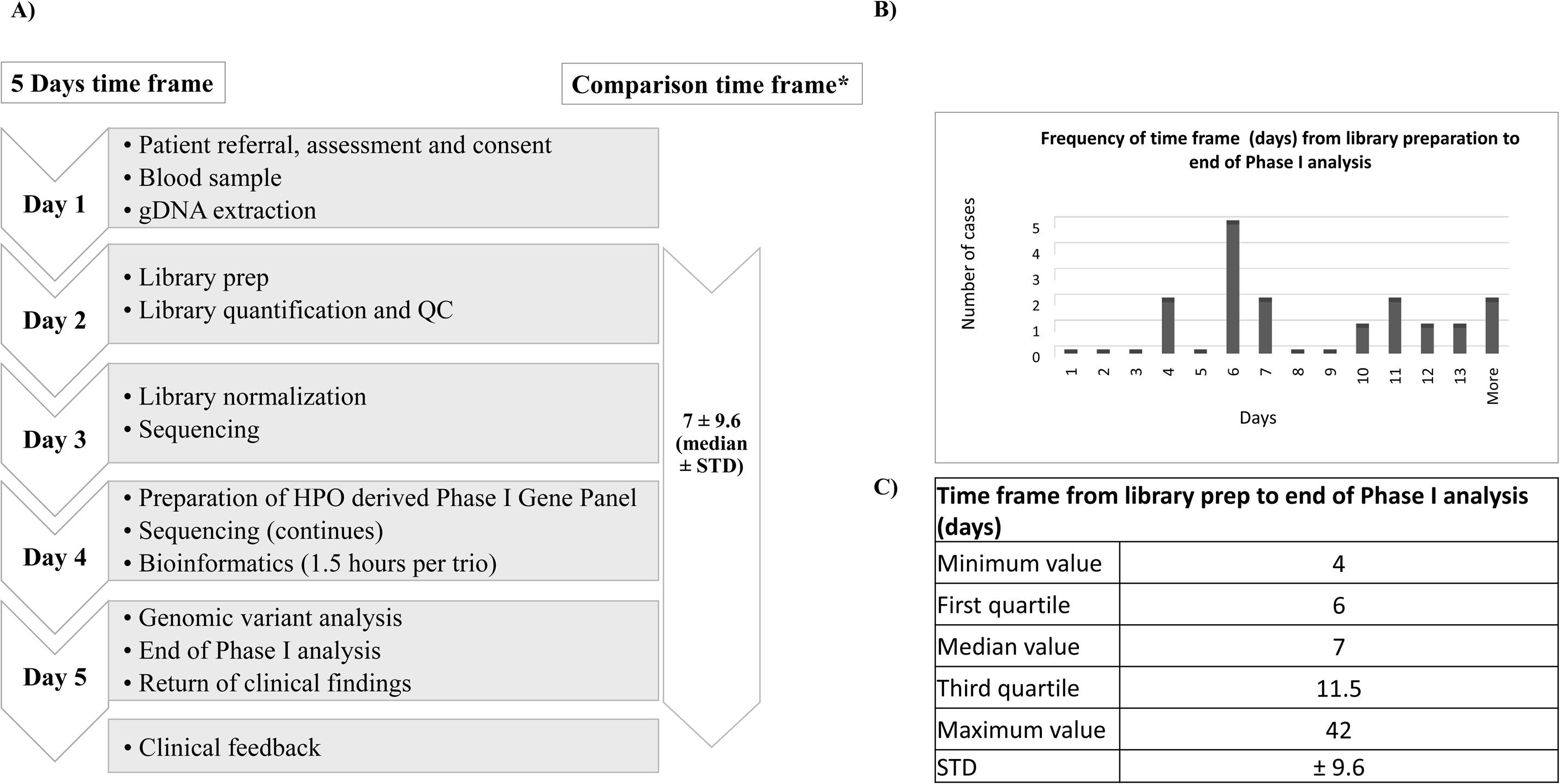
RaPS Time Frame. **RaPS Time Frame for the last 14 cases** (Note, the 10 first cases were used for proof of principle and establishment of the workflow). **A)** The 5 days time frame was achieved as indicated on the left panel. * To provide a comparison with the time frames published by previous studies (13-15) we have calculated the median time frame of the last 14 cases from the time library preparation was initialised. **B)** Histogram of time frame of genomic sequencing calculated from library preparation to return of clinical findings. Weekends, holidays and delays due to reagents failure or unavailability of sequencers are not excluded from the time frame to reflect real life working conditions. **C**) Table shows the quartile distribution of time to diagnosis (calculated from library preparation to return of clinical findings).

### Impact of results on clinical management

In all families where a genetic diagnosis was achieved, diagnosis enabled counselling about prognosis, avoidance of unnecessary investigations and informed recurrence risk. In three individuals (RaPS_02, 11 and 16), a rapid diagnostic result had an immediate impact on the individual’s clinical management.

In the first individual, a molecular diagnosis of *COL3A1* (RaPS_02) associated with vascular EDS helped explain the presence of a ruptured spleen in this individual; prior to this genetic diagnosis child protection concerns had been raised. Secondly, individual RaPS_11, who presented with renal failure was found to have a *de novo WT1* mutation. This genetic diagnosis explained the renal phenotype and also informed the need for bilateral nephrectomy to prevent the development of Wilms tumours that are frequently associated with *WTS1* mutations (26). Finally, the broad approach was especially successful in diagnosing RaPS_16 with Sotos syndrome, an overgrowth disorder (MIM117550). This individual was severely ill with hyperinsulinaemia, and multi-system involvement. The diagnosis of Sotos syndrome was unlikely to have been made for many months in this individual as the clinical features were atypical. A diagnosis of Sotos syndrome assisted in the endocrine management of the hyperinsulinaemia by making further planned investigations unnecessary and advising that this was likely to be self-limiting.

## Discussion

We have developed a robust and readily adoptable protocol for achieving rapid end-to-end WGS-based analysis to support the diagnosis of critically ill children. Our workflow comprises detailed inclusion criteria, clinical data capture using HPO terms, rapid DNA extraction and WGS, a rapid bioinformatics analysis pipeline, and tiered variant reporting.

We have successfully applied this workflow in critically ill children on an intensive care unit in a UK NHS setting and obtained a diagnostic rate of 42% (Table 1). This is a higher diagnostic rate than that obtained in a recently reported study that utilised exome sequencing (9). Furthermore, our time to diagnosis is greatly reduced using WGS, with the shortest time taken to reach a provisional diagnosis being just four days (Figure 3). In three individuals (RaPS_02, 11 and 16) the identification of a diagnostic variant changed the immediate clinical management. In all cases a diagnosis enabled accurate genetic counselling and disease-based management in all families.

Rapid feedback required a close working relationship between the multi-disciplinary teams and for all laboratory and computational systems to be coordinated. A critical part of our workflow was the implementation of phased variant analysis and reporting to facilitate identification of likely causal variants. For successful phased variant reporting in our study, comprehensive phenotypic data captured as HPO terms was required. This enabled rapid generation of appropriate gene panels to clinically assess pathogenicity of variants identified. An example of the clinical utility of a rapid gene panel is highlighted by our analysis of individual RaPS_11, in whom we identified a *WT1* mutation. Typically specific *WT1* testing would take on average eight weeks. However, in this individual the genetic differential diagnosis was wide and in the absence of RaPS WGS, routine genetic testing would have been initiated and likely included a WES gene panel with an expected turnaround time of 4 months. Further, we demonstrate the utility of WGS over WES by the diagnosis made in individual RaPS_24 in whom we identified a compound heterozygous mutation in the gene *CC2D2A* comprising a coding variant and a multi exon-spanning structural variant (inversion) that would not have been identified with WES.

All diagnostic variants, including the previously confirmed *IL2RG* variant in RaPS_08, were identified in Phase I of our tiered reporting system demonstrating its diagnostic utility (Figure 1). We additionally identified a secondary finding of a homozygous *BCHE* variant in Phase II analysis. In this individual a cholinesterase assay confirmed the functional impact of the variant. Although this finding was not relevant to the underlying complex phenotype of this individual it was assessed as important to report back to the clinical team as a secondary finding. This individual underwent several surgical procedures and therefore knowledge that post-anaesthetic apnoea was a risk with certain anaesthetic agents changed their clinical management thus significantly reducing the risk of post-anaesthetic apnoea.

A pivotal part of our data analysis pipeline that enabled rapid diagnosis was the use of fast data processing software for mapping and variant calling (GENALICE, Supplementary material 4). Using this system we were able to significantly reduce the processing time from raw sequence data (FASTQ format) to text files containing lists of variants from the reference sequence (variant call format (VCF) files)(27) from up to 144 hours (using a standard GATK pipeline) to 60 minutes per trio. To ensure the increased processing speed did not adversely affect the accuracy of variant calls we processed the Genome in a Bottle (GIAB) reference sample under exactly the same conditions as our RaPS samples (Supplementary Material 4) (28). Furthermore, the use of Ingenuity Variant Analysis™ software (29) for the annotation and filtering of variants decreased the time taken for interpretation by allowing us to apply our Phased variant analysis models to the data (Figure 1).

It is important to distinguish the difference between our study and that of other groups who have also performed rapid WGS on critically ill children (13-15). In the studies by the Kingsmore group, the time to diagnosis was far quicker owing in part to the manufacturer reconfiguration of the sequencer used and their protocol requiring staff to be available to perform each stage on a non-stop 24 hour cycle. While in the study by van Diemen and colleagues, the analysis of variant data was greatly simplified by analysing a pre-determined list of 3426 genes in all samples. It is also unclear how best to compare our time scales to previous studies as in the calculation of total time they assume no time interval between the various steps of the protocol. Here, we describe a protocol using off-the-shelf reagents and equipment, which fits into the standard working practices of a diagnostic laboratory. We also combine the benefits of a tiered analysis strategy based on a bespoke WGS panel whilst also affording the option of broader unbiased analysis if a diagnosis is not forthcoming.

At present we calculate the cost of running a rapid WGS trio is approximately £5600 but this is likely to fall and needs to be considered in the context of the cost of an ICU bed, estimated to be £4500/day. This cost also compares favourably with the 26 hour protocol previously reported (13) in which reagents were estimated at $6500 per person ($19500 per trio). A full health economics study would be beneficial to extrapolate the health benefits of a rapid diagnosis (versus one taking several months) in this group of severely ill complex individuals as preliminary data suggests sequencing could save up to $7640 per family (30). In the future, all ill patients with suspected genetic disorders will likely have access to WGS. Until then it is important to carefully select those who will benefit most. Given the costs involved in managing critically ill individuals, a rapid genetic diagnosis in this group may ultimately be the most cost effective option for the NHS and other healthcare providers.

In summary, we have presented a sustainable end-to-end workflow for using WGS to rapidly diagnose critically ill individuals with likely monogenic genetic disorders. Such a workflow utilises off-the-shelf products and could readily be adopted by other diagnostic centres.

## Acknowledgements

We gratefully acknowledge the NIHR GOSH BRC funding of the RaPS project. We also extend our gratitude to the many clinicians and scientists at Great Ormond Street Hospital and The Institute of Child Health who have supported the project to date. Thanks especially to the individuals with rare disorders and their families, without whom this work would not have been possible. We thank Professor Lyn Chitty and Dr Stephen Marks for critical review of the manuscript (UCL\GOSH). PB is a NIHR Senior Investigator. All research at Great Ormond Street Hospital NHS Foundation Trust and UCL Great Ormond Street Institute of Child Health is made possible by the NIHR Great Ormond Street Hospital Biomedical Research Centre. The views expressed are those of the author(s) and not necessarily those of the NHS, the NIHR or the Department of Health.

## Contributions

HW, LB, SD, PS and LO performed the laboratory and variant analyses. HW, LB, EC and WJ devised the RaPS workflow and phased variant reporting. PB, CB, HW and LJ initiated and supervised the RaPS workflow. AG and NJ performed the bioinformatics analysis. HW, EC, LB and WJ wrote the manuscript, which was critically reviewed by all authors. EC, WJ, JH, MP and HW liaised with specialist paediatricians to recruit individuals to the study. WJ, EC, RS, SR and JH reviewed variants from Phase I, II and III. Members of the North East Thames Regional Genetics Service and the molecular Genetics laboratory reviewed potentially pathogenic variants at the weekly multidisciplinary team meeting.

## Research in context

### Evidence before this study

We searched PubMed from inception to October 12 2017 using the search terms “whole genome sequencing”, “intensive care” and “human”, for articles published in English. This resulted in 12 publications, however, 8 of these studies corresponded to manuscripts describing microbial or viral pathogen outbreaks in intensive care and so were not relevant. This resulted in only 4 publications relevant to this study and of these 2 describe research finding whilst 2 are perspective articles relating to the original research articles. This is not surprising given it is only recently that whole genome sequencing has been an affordable technique and the novelty of applying such a technique for the clinical diagnosis of critically ill children. The two research articles published in this area are from the same group from the USA where the clinical care environment differs greatly from the publically funded National Health Service (NHS) that operates in the UK. We have in turn sought to establish a framework that is fully compatible to the standard work practices of molecular diagnostic labs throughout the UK and further afield.

### Added value of this study

We have completed a fully prospective study abiding by the standards that are already in place in UK accredited laboratories. Our framework builds on our experiences of integrating the various clinical, diagnostic and research teams who need to come together to make a project such as this a success. We make this information available so that other groups can adopt this approach in their own diagnostic environment. Importantly, all the laboratory and computational equipment we have used is widely available off-the-shelf and the phased variant interpretation algorithm we have used is made open for others to use. We show under real-life conditions that it is possible to obtain a provisional diagnosis within 5 days of identifying a suitable individual and that in a number of cases the identification of a causative gene can have an immediate impact on clinical management decisions.

### Implications of all available evidence

Our study is the first to demonstrate in a UK hospital setting the benefit of using rapid WGS for the diagnosis of critically ill children. The cohort of individuals chosen for this study were selected as they are critically ill and present with a clinically complex phenotype which makes it difficult to make the optimal clinical management decisions. They would clearly benefit from a rapid diagnosis and given the costs involved in their treatment on the paediatric intensive care unit this may represent the most cost effective strategy for the NHS.

