## Supplementary Materials for "Rapid Paediatric Sequencing (RaPS): Comprehensive real-life workflow for rapid diagnosis of critically ill children"

### Slide 1
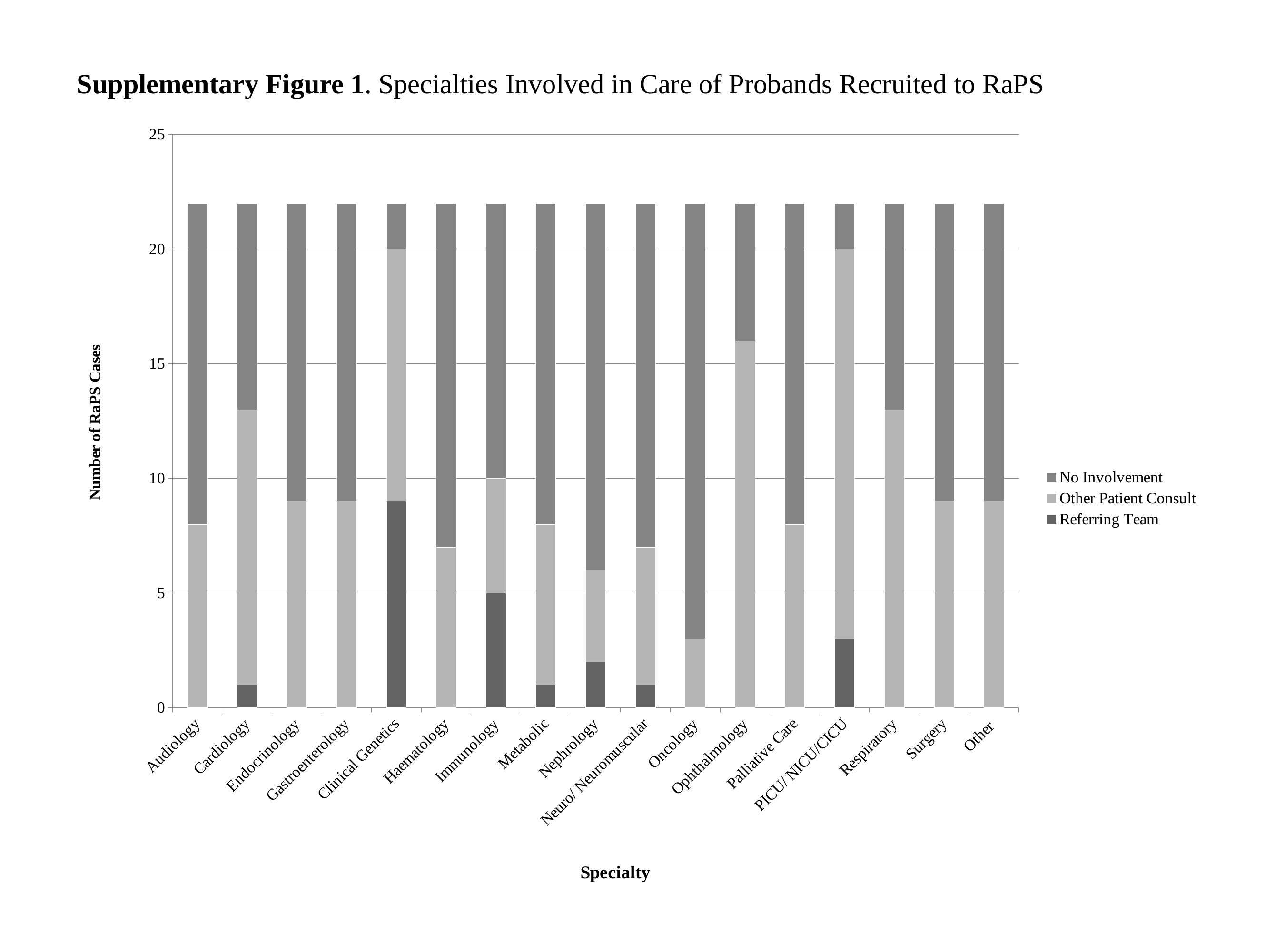

Supplementary Figure 1. Specialties Involved in Care of Probands Recruited to RaPS
#### Chart
| Category | Referring Team | Other Patient Consult | No Involvement |
|---|---|---|---|
| Audiology | 0.0 | 8.0 | 14.0 |
| Cardiology | 1.0 | 12.0 | 9.0 |
| Endocrinology | 0.0 | 9.0 | 13.0 |
| Gastroenterology | 0.0 | 9.0 | 13.0 |
| Clinical Genetics | 9.0 | 11.0 | 2.0 |
| Haematology | 0.0 | 7.0 | 15.0 |
| Immunology | 5.0 | 5.0 | 12.0 |
| Metabolic | 1.0 | 7.0 | 14.0 |
| Nephrology | 2.0 | 4.0 | 16.0 |
| Neuro/ Neuromuscular | 1.0 | 6.0 | 15.0 |
| Oncology | 0.0 | 3.0 | 19.0 |
| Ophthalmology | 0.0 | 16.0 | 6.0 |
| Palliative Care | 0.0 | 8.0 | 14.0 |
| PICU/ NICU/CICU | 3.0 | 17.0 | 2.0 |
| Respiratory | 0.0 | 13.0 | 9.0 |
| Surgery | 0.0 | 9.0 | 13.0 |
| Other | 0.0 | 9.0 | 13.0 |
