## Supplementary Materials for "Rapid Paediatric Sequencing (RaPS): Comprehensive real-life workflow for rapid diagnosis of critically ill children"

### Slide 1
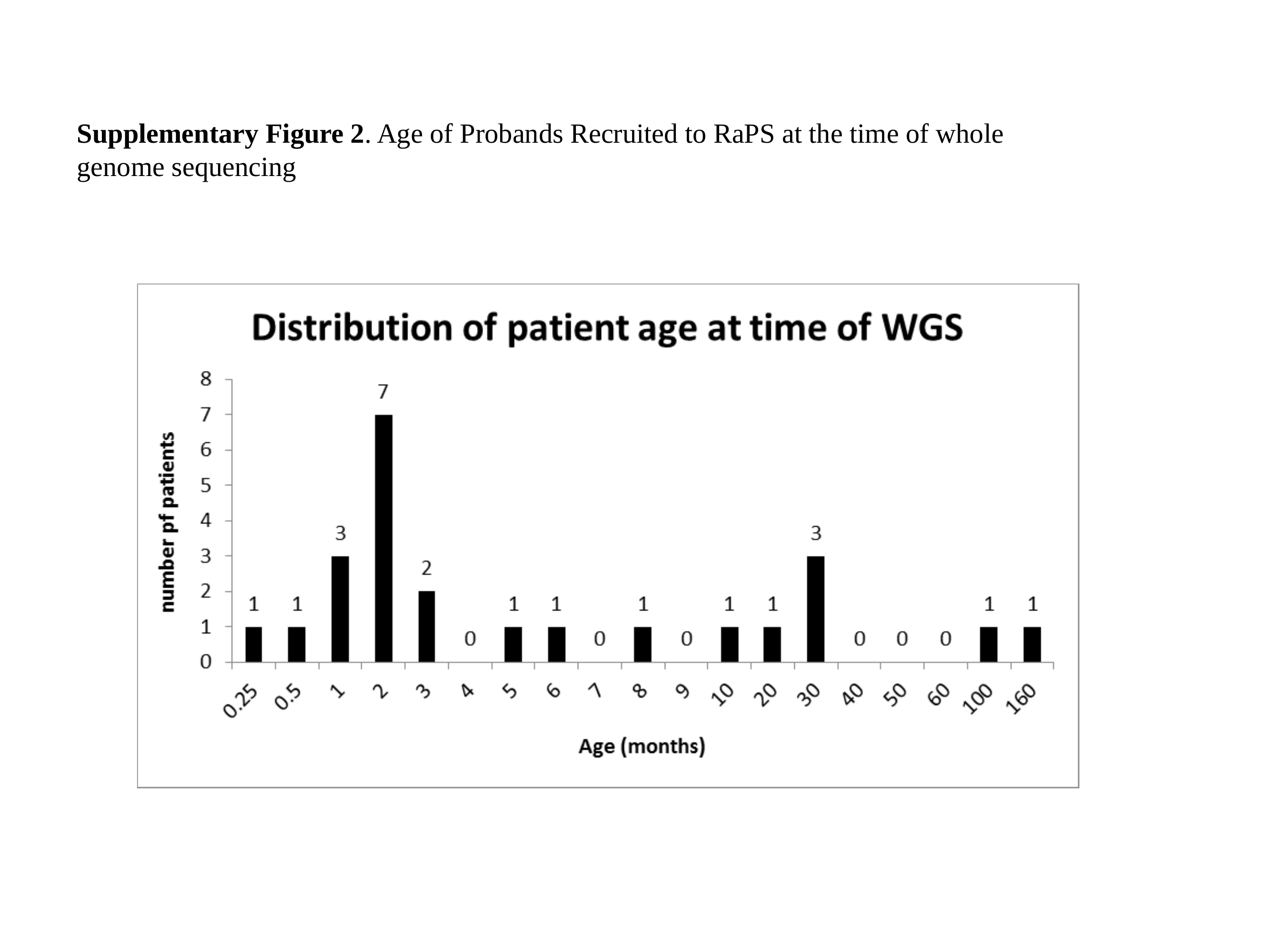

Supplementary Figure 2. Age of Probands Recruited to RaPS at the time of whole genome sequencing
