## Supplementary Materials for "Rapid Paediatric Sequencing (RaPS): Comprehensive real-life workflow for rapid diagnosis of critically ill children"

### Slide 1
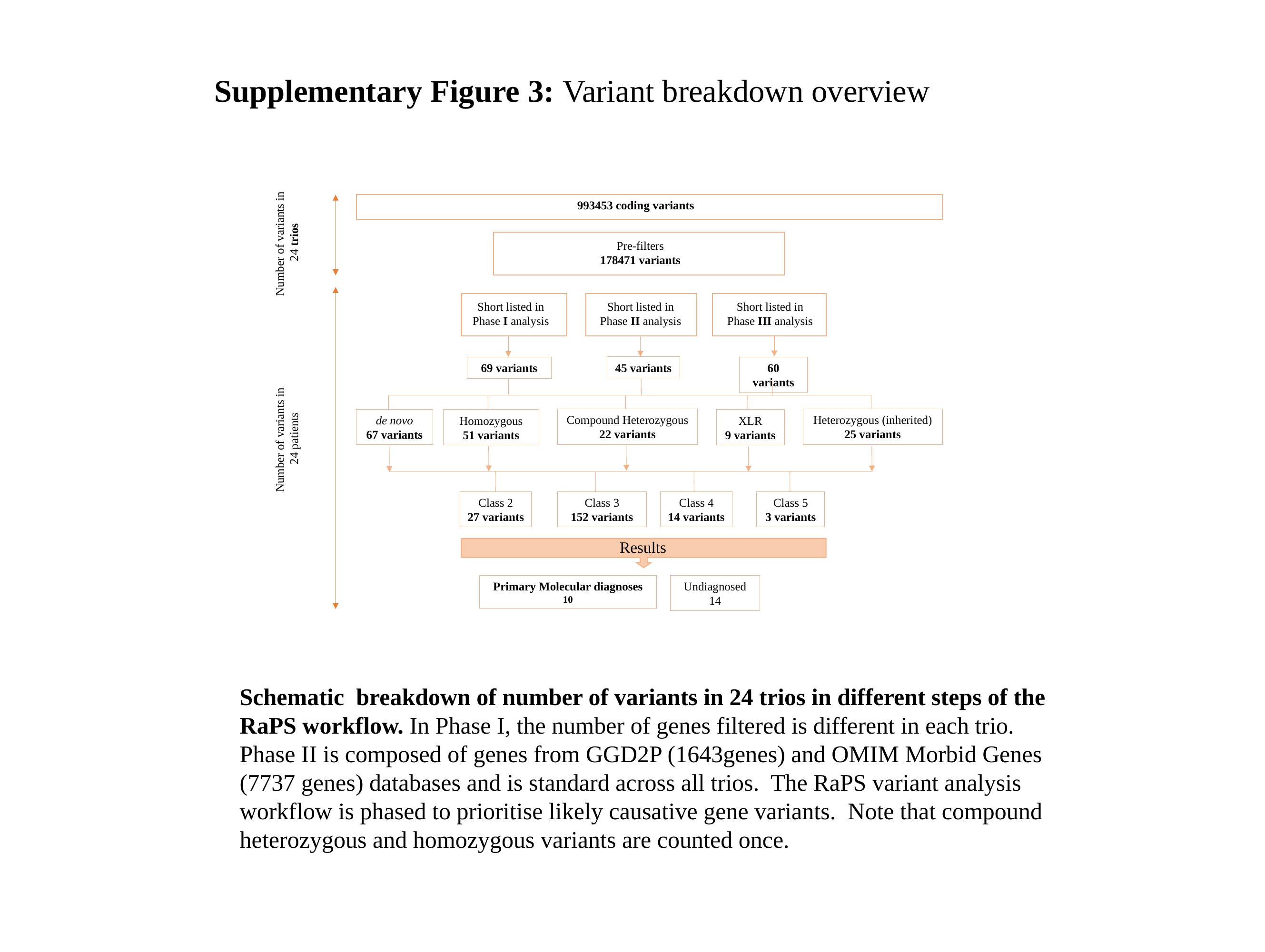

Supplementary Figure 3: Variant breakdown overview
993453 coding variants
Pre-filters
178471 variants
Short listed in Phase II analysis
Short listed in Phase III analysis
Short listed in Phase I analysis
45 variants
69 variants
60 variants
Compound Heterozygous
22 variants
Heterozygous (inherited)
25 variants
de novo
67 variants
XLR
9 variants
Homozygous
51 variants
Class 2
27 variants
Class 3
152 variants
Class 4
14 variants
Class 5
3 variants
Results
Undiagnosed
14
Primary Molecular diagnoses
10
Number of variants in
24 trios
Number of variants in
24 patients
Schematic breakdown of number of variants in 24 trios in different steps of the RaPS workflow. In Phase I, the number of genes filtered is different in each trio. Phase II is composed of genes from GGD2P (1643genes) and OMIM Morbid Genes (7737 genes) databases and is standard across all trios. The RaPS variant analysis workflow is phased to prioritise likely causative gene variants. Note that compound heterozygous and homozygous variants are counted once.
